# Wobbly hedgehog syndrome- a progressive neurodegenerative disease

**DOI:** 10.1101/2023.07.13.547983

**Authors:** Grayson A. Doss, Daniel Z. Radecki, Arya Kethireddy, Madelyn J. Reilly, Andrea E. Pohly, Benjamin K. August, Ian D. Duncan, Jayshree Samanta

**Affiliations:** Department of Surgical Sciences, School of Veterinary Medicine, University of Wisconsin-Madison, 2015 Linden Drive, Madison, WI 53706, USA; Department of Comparative Biosciences, School of Veterinary Medicine, University of Wisconsin-Madison, 2015 Linden Drive, Madison, WI 53706, USA; Department of Pathobiological Sciences, School of Veterinary Medicine, University of Wisconsin-Madison, 2015 Linden Drive, Madison, WI 53706, USA; University of Wisconsin-Madison School of Medicine and Public Health Electron Microscope Facility; Department of Medical Sciences, School of Veterinary Medicine, University of Wisconsin-Madison, 2015 Linden Drive, Madison, WI 53706, USA; Current address: Department of Biomedical Sciences, College of Veterinary Medicine, University of Georgia, 501 DW Brooks Drive, Athens, GA 30602, USA

**Keywords:** Neurodegeneration, progressive multiple sclerosis, remyelination, demyelination, neuronal cell death, aberrant myelin.

## Abstract

Wobbly hedgehog syndrome (WHS) has been long considered to be a myelin disease primarily affecting the four-toed hedgehog. In this study, we have shown for the first time that demyelination is accompanied by extensive remyelination in WHS. However, remyelination is not enough to compensate for the axonal degeneration and neuronal loss, resulting in a progressive neurodegenerative disease reminiscent of progressive forms of multiple sclerosis (MS) in humans. Thus, understanding the pathological features of WHS may shed light on the disease progression in progressive MS and ultimately help to develop therapeutic strategies for both diseases.

**Highlights:**

1. Wobbly hedgehog syndrome (WHS) is a progressive neurodegenerative disease.
2. Spongy degeneration of the brain and spinal cord is the diagnostic feature of WHS.
3. WHS affected brain and spinal cord show extensive demyelination and remyelination.
4. Axonal degeneration is accompanied by loss of neurons in WHS.

Graphical Abstract

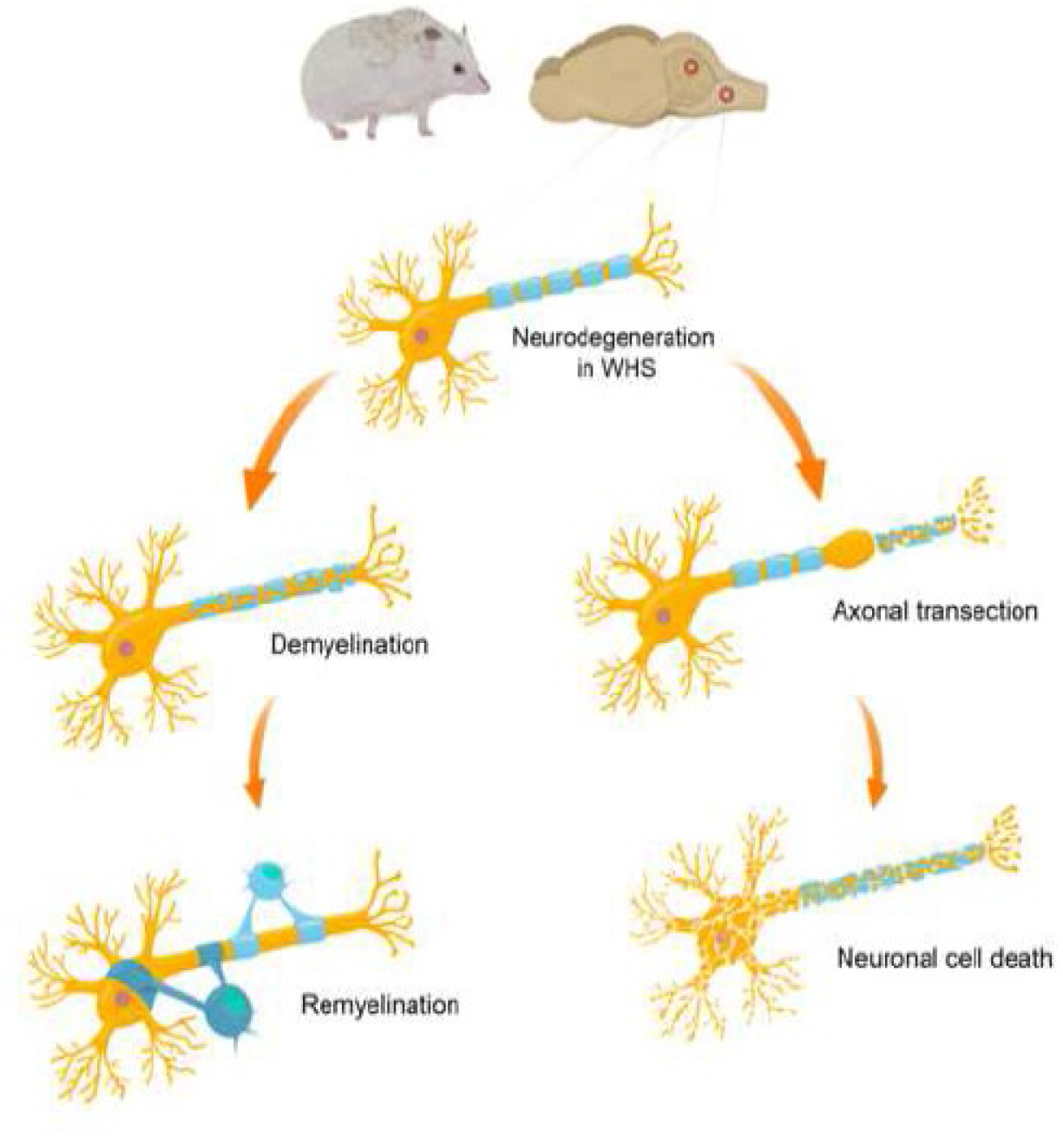

## 1. Introduction

Wobbly hedgehog syndrome (WHS) is a progressive fatal neurological disease of four-toed hedgehogs (*Atelerix albiventris*), also known as African pygmy hedgehogs, a popular companion animal. It is characterized by vacuolation of the white matter of the brain and spinal cord, with the majority of lesions being found in the cerebellum, brainstem, and the cervical and thoracic spinal cord (Diaz-Delgado et al., 2018). Affected hedgehogs usually present with ataxia and weakness in both hindlimbs (paraparesis) at around two years of age, but the disease has also been reported in younger animals. The weakness rapidly deteriorates to involve all four limbs (tetraparesis) resulting in loss of ambulation with inability to walk and stand upright, and ultimately progressing to death (Graesser et al., 2006). Although the prevalence of WHS is suspected to be high, with reports estimating that up to 10% of captive hedgehogs in North America may be affected, the etiology of WHS remains unknown (Graesser et al., 2006). A similar disorder has been reported in an outbreak in European hedgehogs (*Erinaceus europaeus*), though it appears to have been self-limiting (Palmer et al., 1998).

Based on the presence of vacuolation of the white matter, WHS is assumed to be a primary myelin disorder. Myelin consists of spiral wraps of oligodendrocyte processes around axons and contains about 70% lipids. Its main function is to insulate the axons for faster conduction of action potentials, thus increasing the nerve conduction velocity. Myelin can be lost primarily due to loss of oligodendrocytes or secondarily due to loss to neurons. Loss of myelin occurs in many neurodegenerative diseases especially multiple sclerosis (MS), an autoimmune myelin disease in humans. The predominantly late age of onset and progressive clinical course of WHS is very similar to primary and secondary progressive forms of MS (Graves et al., 2022). However, the inflammatory reaction is less prominent in progressive MS compared to the most common relapsing remitting form of MS (Frischer et al., 2009; Revesz et al., 1994; Trapp and Nave, 2008). Consequently, disease modifying drugs that target the immune system are less effective for treatment of progressive MS (Amezcua, 2022). Interestingly, treatment with immune modulators like steroids and interferon-β1a, approved for therapy of MS, has been attempted in WHS affected hedgehogs but has failed to alter the clinical course of the disease (Graesser et al., 2006). Another prominent feature of long-standing, progressive MS, is atrophy of the brain resulting from extensive neuronal loss due to failure of functional compensation by remyelination (Lassmann et al., 2012; Trapp and Nave, 2008). However, definitive evidence of demyelination and remyelination, axonal degeneration, and neuronal loss has not been reported in WHS affected hedgehogs.

The current knowledge of WHS has been largely gained from individual case reports with very little comparison of the findings to healthy hedgehogs (Diaz-Delgado et al., 2018; Graesser et al., 2006; Madarame et al., 2014; Palmer et al., 1998; Silva et al., 2022). In this study, we compared the brain and spinal cord of multiple four-toed hedgehogs diagnosed with WHS to similar numbers of age and sex matched healthy animals. Specifically, we examined the myelin and neuronal pathology in client-owned hedgehogs affected by WHS euthanized at late stages of the disease and compared them to hedgehogs without neurological signs. Our results show that extensive demyelination and remyelination occur along with widespread cell death of neurons in the cerebellum and spinal cord of WHS affected hedgehogs. Contrary to the prevailing view, our study indicates that WHS is a progressive neurodegenerative disease in which remyelination is present but ongoing myelin vacuolation and neuronal apoptosis leads to chronic and terminal disease.

## 2. Material and methods

### 2.1. Animal care

This study was approved by the University of Wisconsin-Madison School of Veterinary Medicine Institutional Animal Care and Use Committee. Affected hedgehogs consisted of captive-bred animals with end-stage neurologic disease enrolled upon owner consent. Control animals were captive bred hedgehogs without neurological signs. Hedgehogs received a full neurological examination consisting of awake evaluation of behavior, gait, mentation, as well as selected cranial nerves and reflexes (Berg et al., 2021).

### 2.2. Sample collection and processing

Animals were anesthetized using vaporous isoflurane until a surgical plane of anesthesia was reached based on loss of the pelvic limb withdrawal reflex. The animals were perfused with intracardiac administration of 1mL of heparin sulfate to prevent the blood from clotting followed by 180 mL of phosphate buffered saline (PBS) at 20-30 mL/min to drain the blood and finally with 180 mL of 4% PFA in PBS at 20-30 mL/min using a perfusion pump. The brain, cervical spinal cord, and middle ear were harvested from the perfused animals. The middle ear was stored in 10% neutral buffered formalin until analysis. The brains were post-fixed in 4% paraformaldehyde (PFA) for 2-3 hours at room temperature followed by incubation in 30% sucrose solution at 4ᴼC on a shaker for 12-18 hours. The brains were then cryoprotected by embedding in Tissue-Tek O.C.T. (Sakura, #4583) and frozen at –80ᴼC for at least 24 hours before sectioning (Clawson et al., 2023). A Leica CM3050S cryostat was used to obtain 20 µm sagittal sections of the brains and the slides were stored at –80ᴼC until further analysis.

### 2.3. Luxol fast blue – hematoxylin and eosin staining

The brain cryosections were first stained with luxol fast blue (LFB) with minor modifications (Carriel et al., 2017). Briefly, LFB solution was prepared by dissolving Solvent Blue-38 (Sigma #S3382) powder in 80% ethanol to create a 1% w/v solution. The slides were washed in increasing concentrations of ethanol and placed in LFB at 65-70ᴼC for 12-16 hours, protected from light. Slides were then washed in 80% ethanol for 2-3 minutes and differentiated in 0.1% lithium carbonate (Sigma #62470) for 45-60 seconds followed quickly by washes in distilled water to stop the differentiation. Following LFB staining, slides were stained with hematoxylin and eosin (H&E) using the standard protocol (Feldman and Wolfe, 2014).

### 2.4. Immunofluorescence

The brain cryosections were immunolabeled with mouse anti –Olig2, –CC1, –MOG, –βIII-tubulin, rabbit anti –HuC/D, –cleaved caspase 3, and rat anti-PLP1 (Table S1) using published protocols (Clawson et al., 2023).

For analysis of CC1 fluorescence intensity, we used the Hybrid Cell Counter function of Keyence Image Analysis software (Keyence Corp. of America, Elmwood, New Jersey) and quantified all the CC1 positive cells with area >10 µm^2^ and a fluorescence intensity >25 A.U., containing a Hoechst labelled nucleus, without smoothing the image.

### 2.5. Ultrastructural analysis

Following intracardiac perfusion with 4% PFA, the cervical spinal cord was dissected and immersion post-fixed in 2.5% glutaraldehyde for at least 48 hours. Spinal cord sections were trimmed and embedded in plastic molds. Images were acquired using Philips CM120 transmission electron microscope (TEM). Axon diameter, number of axons, and g-ratio were calculated using the semi-automated MyelTracer program (Kaiser et al., 2021). Briefly, axon outlines were defined using the thresholding tool and manually traced to remove edge effects, followed by threshold identification of the adaxonal myelin membrane (inner edge). Next, the abaxonal myelin membrane (outer edge) was delineated manually and any axons with myelin pathologies or artifacts (e.g., delamination, swelling etc.) were excluded from g-ratio calculations but included in total axon counts.

### 2.6. Statistical analysis

All experiments except ultrastructural analysis were replicated at least three times. For immunofluorescent experiments, at least five sections per animal were analyzed and data from three to five animals were combined. For ultrastructural analysis, more than 500 axons were counted from multiple areas of the cervical spinal cord of one animal per group (i.e., healthy, aged WHS affected, and young WHS affected hedgehogs). All the data are expressed as mean ± standard error of mean in the graphs and mean ± standard deviation in the results. Statistical analysis was performed using Student’s t test, or two-way ANOVA with Tukey’s post-hoc t test. Differences were considered statistically significant at p < 0.05.

## 3. Results

### Histopathological diagnosis of WHS

The diagnostic criteria for WHS includes vacuolation or spongy degeneration of the brain and spinal cord associated with a prior clinical history of ataxia and progressive ascending paralysis (Diaz-Delgado et al., 2018; Graesser et al., 2006; Madarame et al., 2014; Palmer et al., 1998; Silva et al., 2022). To confirm the pathological diagnosis of WHS, we performed histochemical analysis of the cerebellum, brainstem, and cervical spinal cord from suspected WHS hedgehogs to that of age and sex matched hedgehogs without neurological signs. As previously reported, the predominant change in the white matter in WHS was myelin vacuolation. In the cerebellar white matter, LFB staining showed widespread vacuolation along with faint staining in the areas surrounding the vacuoles (Fig. 1A), similar to prior reports (Diaz-Delgado et al., 2018; Silva et al., 2022). These areas of faint LFB staining have been described as shadow plaques in the brains of people with MS and are considered a hallmark of remyelination (Franklin, 2002). Thus, our results point to the possibility of remyelination triggered by demyelination in the vacuolated areas. These observations were confirmed in 1 micron toluidine blue stained sections of the affected cervical spinal cords which showed a range from multiple focal areas of vacuolated fibers (Fig. 1B) to scattered focal areas, primarily in the lateral and ventral columns (Fig. S1).

**Figure 1:**
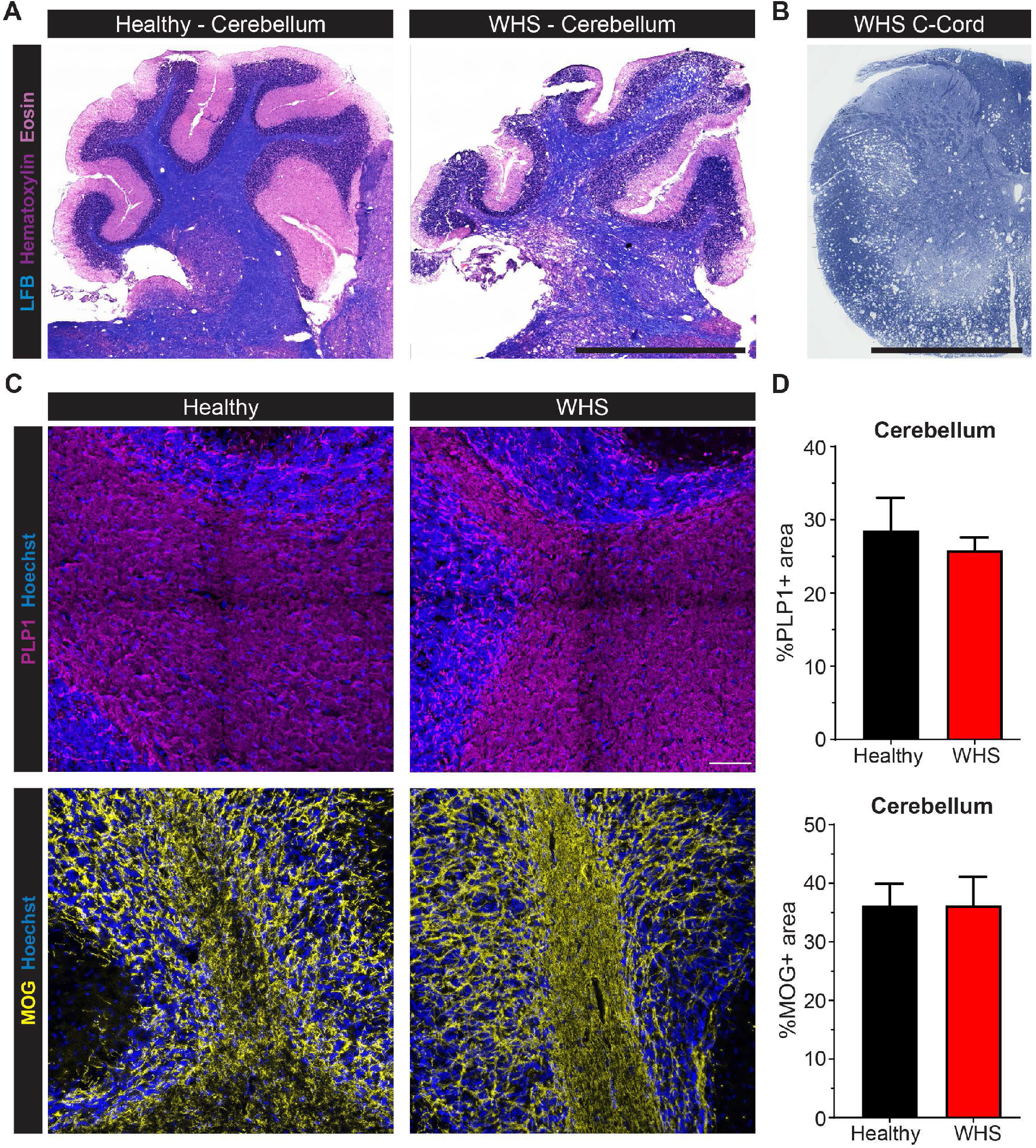
Histopathological diagnosis of WHS. (A) Luxol fast blue (LFB) with hematoxylin & eosin staining of cerebellum shows vacuolation and faint staining in WHS affected hedgehog, n = 3 hedgehogs/group, Scale = 25 µm. (B) Toluidine blue staining of WHS affected cervical spinal cord shows extensive vacuolation in the lateral and ventral column, n = 3, Scale = 25 µm. (C) Immunofluorescent labeling of early myelin protein PLP1 (magenta, top), late myelin protein MOG (yellow, bottom), and nuclei with Hoechst (blue) in the cerebellar white matter, n = 3 hedgehogs/group, Scale = 100 µm. (D) Quantification of PLP (top) and MOG (bottom) positive areas show no difference between the healthy and WHS affected cerebellar white matter, n = 3 hedgehogs/group, student’s t-test.

To further examine the myelin abnormalities, we performed immunofluorescent labelling for myelin proteins with Proteolipid protein (PLP), whose expression begins early in the premyelinating stage of oligodendrocytes, and Myelin Oligodendrocyte Glycoprotein (MOG), which is expressed later in mature oligodendrocytes (Grinspan et al., 1993; Solly et al., 1996) (Fig. 1C). Surprisingly, the area of PLP expression (25.8 ± 1.8% in WHS vs 28.5 ± 4.5% in healthy) and MOG expression (36.2 ± 4.9% in WHS vs 36.2 ± 3.7% in healthy) were similar in the cerebellum of affected and healthy hedgehogs (Fig. 1D). Overall, these results suggest that the myelin proteins are replenished by remyelination in the cerebellum of the WHS affected hedgehogs.

Finally, to rule out middle ear disease as a cause of ataxia, we examined the bullae of the affected hedgehogs with H&E staining and did not detect any abnormalities (Fig. S2). Thus, central nervous system disease is the cause of clinical signs in WHS affected hedgehogs.

### Evidence of demyelination and remyelination in WHS

Ultrastructural examination of the spinal cord from a young (6.5 months) and an older (26 months) animal with WHS, showed the presence of vacuolation along with evidence of widespread demyelination and remyelination. Demyelinated axons were seen singly or in small groups in both ages with ∼13-fold higher number of demyelinated axons in the WHS affected spinal cord (41.02 ± 26.02%) compared to healthy spinal cord (3.39 ± 3.11%) (Fig. 2A,D). Axons with thin myelin sheaths indicative of remyelination, were seen scattered throughout the neuropil and in small groups (Fig. 2A, Fig. S3). To confirm that these were remyelinated, we calculated the g-ratio, which is the ratio of axon diameter to the fiber diameter (Duncan et al., 2017). The g-ratio of axons in the affected hedgehogs was significantly higher (0.75 ± 0.94, 247 axons) compared to that in the healthy hedgehogs (0.727 ± 0.078, 414 axons) (Fig. 2B). In particular, axons with larger diameters (i.e., more than 1.5 µm) had higher g-ratios suggesting thinner myelin sheaths observed in remyelinated axons (Fig. 2C). In the younger affected hedgehog, axons with thin myelin sheaths were present in certain tracts such as the sub-pial dorsolateral column (Fig. S4). Deeper to these superficial areas and adjacent to the grey matter, axons were normally myelinated. In contrast, some axons in the sub-pial contralateral tract also had thin myelin but this was less extensive (Fig. S4). These observations demonstrate that the thin myelin sheaths do not result from a developmental hypomyelination but from remyelination. In the older affected hedgehog, axons with thin myelin sheaths were present throughout the cervical spinal cord. Remyelination was also seen in 1 µm sections from three other affected hedgehogs, confirming that myelin repair is a uniform feature of WHS. Despite the extensive vacuolation and demyelination in severely affected hedgehogs, there were many surviving axons (Fig. S5) in addition to axons showing accumulation of organelles suggesting degeneration (Fig. S6).

**Figure 2:**
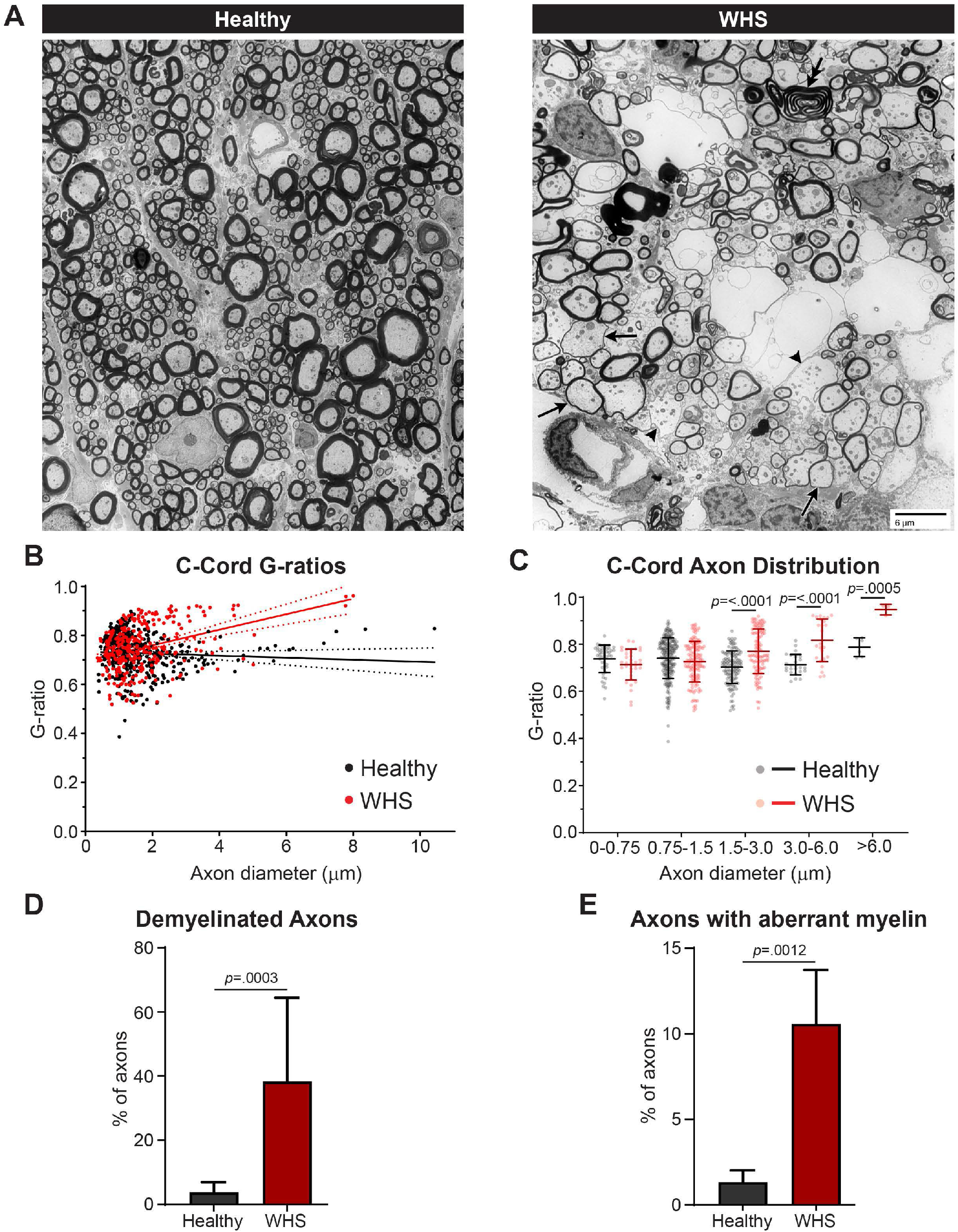
Demyelination and remyelination in the spinal cord in WHS. (A) TEM images from the ventral column of a healthy and WHS affected spinal cord. In the affected animal, multiple vacuoles and a few intact, normally myelinated axons were present. There were numerous demyelinated (arrow heads, twenty-one total), and thinly myelinated axons (arrows, 59 total). A single aberrant myelin sheath was also seen (double arrow). Scale = 6 µm. (B) G-ratio of individual axons in the lateral and ventral columns of cervical spinal cord showed an increase in g-ratio in WHS affected axons >2 µm in diameter (average g-ratio: healthy = 0.727 ± 0.078, axons = 414; WHS = 0.750 ± 0.094, axons = 247). (C) Distribution of g-ratio showed an increase in axon diameter groups–1.5-3 µm, 3-6 µm, and >6 µm. Multiple unpaired t-tests with Welch correction, and Holm-Sidak multiple comparison correction. (D) Quantification of number of demyelinated axons in the spinal cords of healthy (1445 total axons) and WHS affected (1153 total axons) animals, student’s t-test. (E) Quantification of number of axons with aberrant myelin in the spinal cords of healthy (1445 total axons) and WHS affected (1153 total axons) animals, student’s t-test.

A common observation in areas of vacuolation and remyelination was redundant myelin loops/sheaths along with myelin blebbing, outfoldings or protrusions and myelin degeneration (Fig. 2A, Fig. S7). The affected hedgehogs had a 5-fold higher number of axons with aberrant myelin (10.58 ± 3.15%) than in the healthy cervical spinal cord (1.33 ± 0.68%) (Fig. 2E). These aberrant myelin features have been described both during development by newly generated oligodendrocytes and during aging by degenerating oligodendrocytes (Djannatian et al., 2023; Peters, 2002). However, we observed similar pathological myelin features in the younger hedgehog affected with WHS, suggesting that the changes in myelin sheaths are not likely to be due to aging.

Remyelination can be achieved by generation of new mature oligodendrocytes from oligodendrocyte precursor cells (OPCs) (Chapman et al., 2023; Franklin and Ffrench-Constant, 2017; Snaidero et al., 2020) or by extension of new myelin processes from surviving old oligodendrocytes (Bacmeister et al., 2022; Duncan et al., 2018; Yeung et al., 2019). However, the myelin produced by surviving older oligodendrocytes is often mistargeted to the cell bodies of neurons (Franklin et al., 2021; Neely et al., 2022). Accordingly, we observed significantly higher numbers of HuC/D labelled neuronal cell bodies wrapped by PLP positive myelin in the affected cerebellum (6106 ± 1875/mm^3^) compared to the healthy cerebellum (1238 ± 247.5/mm^3^) (Fig. 3). Taken together, the ultrastructural and immunofluorescence data confirms the presence of ongoing remyelination in WHS affected hedgehogs.

**Figure 3:**
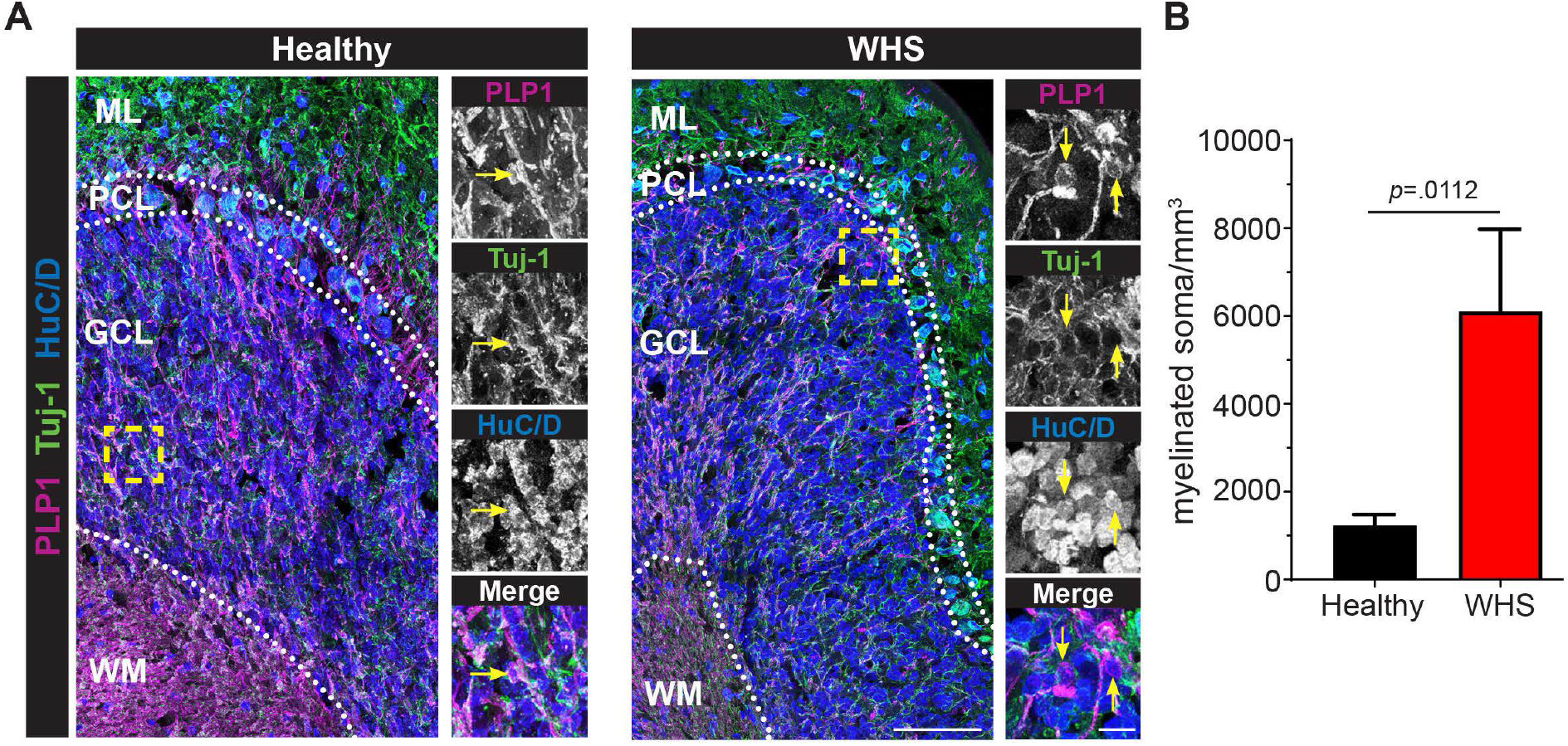
Increase in myelin mistargeting in WHS cerebellum. (A) Immunofluorescent labeling of myelin with PLP1 (magenta), neuronal processes with βIII-tubulin/Tuj-1 (green), and neuronal soma with HuC/D (blue) in the healthy and WHS affected cerebellum show PLP1+HuC/D+ myelinated neuron cell bodies (insets, yellow arrows). WM = white matter; GCL = granule cell layer; PCL = Purkinje cell layer; ML = molecular layer, scale= 100 µm, inset = 10 µm. (B) Quantification of HuC/D+ neuronal soma wrapped by PLP1 show an increase in aberrantly myelinated neuronal soma in the HWS affected cerebellum, n = 3 hedgehogs/group, 2-tailed unpaired t-test.

### Regeneration of oligodendrocytes in WHS

To assess regeneration of oligodendrocytes, we first quantified the number of mature oligodendrocytes labelled with CC1 antibody in the cerebellar white matter (Fig. 4A). Surprisingly, we did not find any difference in the density of oligodendrocytes between healthy (35.4 ± 0.57 X10^3^/mm^3^) and affected hedgehogs (34.05 ± 7.85 X10^3^/mm^3^) (Fig. 4B). However, CC1 labelled oligodendrocytes in the affected cerebellum appeared brighter and labelled a larger area of the soma. CC1 antibody detects the RNA binding protein Quaking 7, which is highly upregulated when pre-myelinating oligodendrocytes transition into myelinating oligodendrocytes (Bin et al., 2016; Wu et al., 2001). Hence, we quantified the number of CC1+ oligodendrocytes with a fluorescence intensity above 25 AU (arbitrary units) and CC1 labelled area >10 µm^2^. The WHS affected cerebellar white matter had significantly higher number of oligodendrocytes above threshold intensity and labelled area compared to the healthy hedgehogs (affected: 1296 ± 530.7 X10^3^/mm^3^ vs healthy: 356.7 ± 141.5 X10^3^/mm^3^) (Fig. 4C,D), indicating regeneration of oligodendrocytes in the affected hedgehogs. However, the total number of cells in the oligodendroglial lineage labelled by Olig2 remained the same in the healthy (170.3 ± 19.01 X10^3^/mm^3^) and affected cerebellum (173.7 ± 70.47 X10^3^/mm^3^) (Fig. 4 E,F). Since the antibodies used to label oligodendrocyte precursor cells (OPCs), NG2 and PDGFRα, failed in hedgehog tissue, we examined the number of OPCs indirectly by quantifying the number of Olig2 labelled cells that did not express the mature oligodendrocyte marker CC1. However, we did not observe a difference between the number of OPCs in the cerebellum of WHS affected hedgehogs (39.37 ± 12.04 X10^3^/mm^3^) and healthy hedgehogs (25.43 ± 5.47 X10^3^/mm^3^) (Fig. 4G). Together, these results suggest that WHS affected hedgehogs not only have a larger proportion of remyelinating oligodendrocytes but also regeneration of oligodendrocytes and OPCs likely compensates for their loss. To understand whether the regeneration was in response to oligodendroglial cell death, we quantified the Olig2 labelled cells co-expressing the apoptosis marker cleaved caspase 3, but did not find any difference in the number of apoptotic cells in the WHS affected (3.77 ± 2.11 X10^3^/mm^3^) versus healthy (7.17 ± 1.02 X10^3^/mm^3^) cerebellar white matter (Fig. 4H,I). Thus, apoptosis of oligodendrocytes is not likely to be the primary cause of remyelination in late stages of the disease. However, we were unable to examine apoptosis in the early stages of the disease, since the WHS affected hedgehogs were client-owned animals euthanized at the late stage of the disease, when the quality of life became poor. Thus, we cannot rule out oligodendroglial cell death triggering remyelination earlier in the disease course.

**Figure 4:**
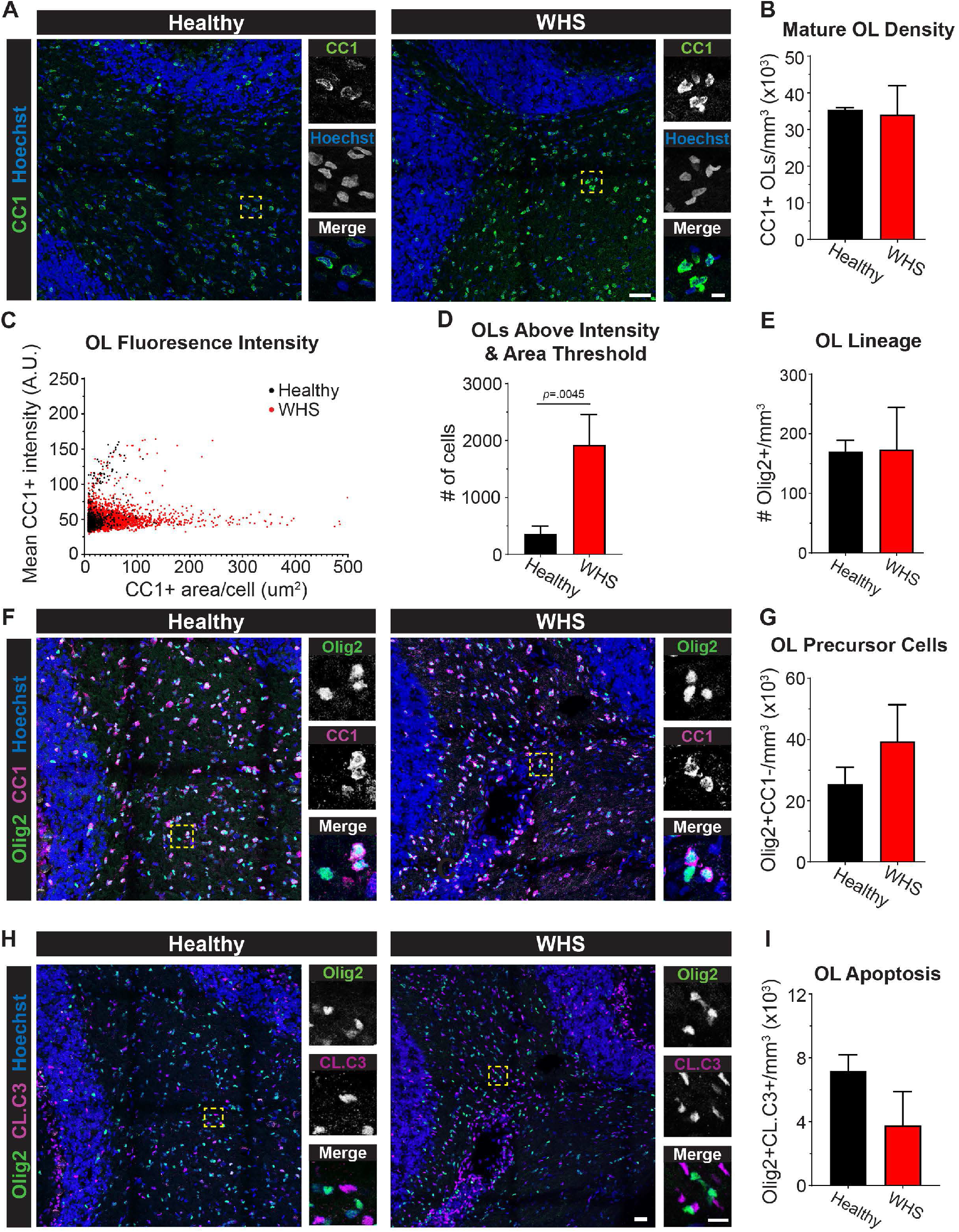
Evidence of newly generated oligodendrocytes in WHS cerebellum. (A) Immunofluorescent labeling of mature oligodendrocyte (OL) with CC1 (green) and nuclei with Hoechst (blue) show brighter and larger CC1 labelled OL soma in WHS affected cerebellum, n = 3 hedgehogs/group, scale = 100 µm, inset = 10 µm. (B) Quantification of total number of CC1+ oligodendrocytes in the cerebellum show no difference between healthy and affected animals, n = 3 hedgehogs/group, 2-tailed t-test. (C) Dot plot showing the cells with CC1 fluorescence intensity above 25 AU and with CC1 labelled soma >10 µm^2^ area. (D) Quantification of OLs with CC1 fluorescence intensity above 25 AU and with CC1 labelled soma >10 µm^2^ area show higher number of cells in WHS affected cerebellum, n = 3 hedgehogs/group, 2-tailed t-test. (E) Quantification of total number of cells in the oligodendroglial lineage labelled with Olig2 show no change between the healthy and WHS affected cerebellum, n = 3 hedgehogs/group, 2-tailed t-test. (F) Immunofluorescent images of oligodendroglial lineage cells labelled with Olig2 (green), mature OLs labelled with CC1 (magenta), and nuclei labelled with Hoechst (blue) in the healthy and WHS affected cerebellum, scale= 50 µm, inset = 10 µm. (G) Quantification of oligodendrocyte precursor cells (Olig2 positive and CC1 negative) show no difference between the healthy and WHS affected cerebellum, n = 3 hedgehogs/group, 2-tailed t-test. (H) Immunofluorescent images of oligodendroglial lineage cells labelled with Olig2 (green), apoptotic cells labelled with cleaved caspase 3 (CL.C3) (magenta), and nuclei labelled with Hoechst (blue) in the healthy and WHS affected cerebellum, scale = 50 µm, inset = 10 µm. (**I**) Quantification of apoptotic oligodendroglial cells (Olig2 and CL.C3 double positive) show no difference between the healthy and WHS affected cerebellum, n = 3 hedgehogs/group, 2-tailed t-test.

### Neuronal cell death is a major driver of progressive disease in WHS

The progressive clinical course of WHS suggests ongoing degeneration in the brain and spinal cord. Therefore, we examined neuronal cell death by quantifying the number of HuC/D labelled neurons co-expressing cleaved caspase 3 in the cerebellum (Fig. 5A). Interestingly, the granule cell layer of the cerebellum in WHS affected hedgehogs showed significantly higher number of apoptotic neurons (affected: 40.33 ± 7.39 X10^3^/mm^3^ vs healthy: 13.37 ± 11.53 X10^3^/mm^3^) (Fig. 5B). In addition, we observed signs of axonal degeneration including axonal spheroids surrounded by myelin in the granule cell layer. But their occurrence was not significantly different between WHS affected (5445 ± 2158 /mm^3^) and healthy (3465 ± 495 /mm^3^) cerebellum (Fig. 5C,D), suggesting that these changes are likely a result of aging rather than active axonal degeneration (Nikic et al., 2011; Trapp et al., 1998). Similarly, we observed occasional axons with accumulation of organelles suggesting early axon degeneration, but surviving axons were present in many, though not all vacuoles, in the affected spinal cords when evaluated with TEM (Fig. S6).

**Figure 5:**
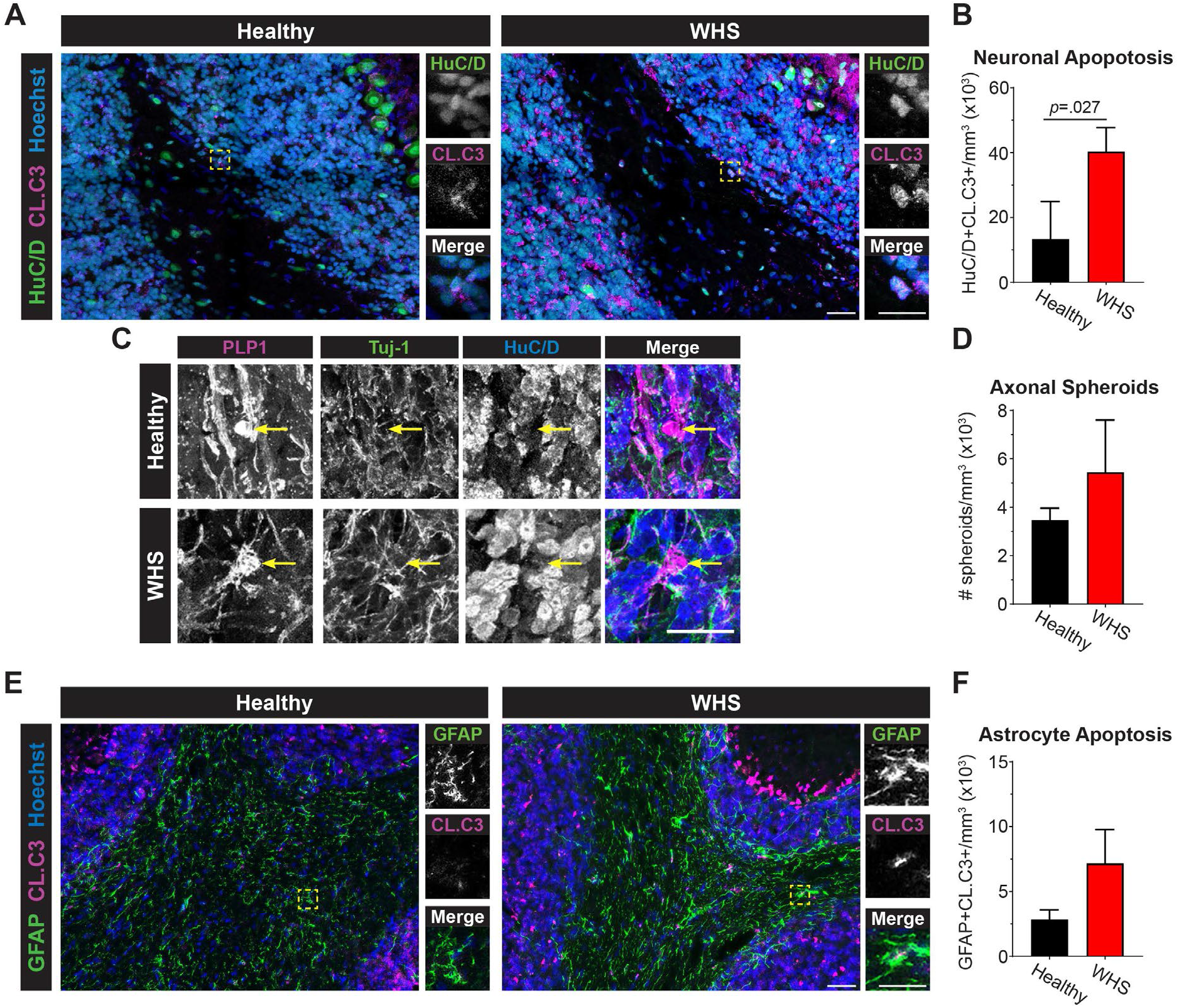
Increase in neuronal apoptosis in WHS affected cerebellum. (A) Immunofluorescent images of neurons labelled with HuC/D (green), apoptotic cells labelled with cleaved caspase 3 (CL.C3) (magenta), and nuclei labelled with Hoechst (blue) in the healthy and WHS affected cerebellum, scale = 30 µm, inset = 15 µm. (B) Quantification of apoptotic neurons (HuC/D and CL.C3 double positive) show increase in apoptosis in the granule cell layer of WHS affected cerebellum, n = 3 hedgehogs/group, 2-tailed t-test. (C) Immunofluorescent images of neuronal processes labelled with Tuj-1 (green), neuronal soma labelled with HuC/D (blue), and myelin labelled with PLP1 (magenta) in the healthy and WHS affected cerebellum, scale = 10 µm. (D) Quantification of axonal spheroids (HuC/D negative and Tuj-1 positive) wrapped with PLP1+ myelin show no change in the granule cell layer of healthy and WHS affected cerebellum, n = 3 hedgehogs/group, 2-tailed t-test. (E) Immunofluorescent images of astrocytes labelled with GFAP (green), apoptotic cells labelled with cleaved caspase 3 (CL.C3) (magenta), and nuclei labelled with Hoechst (blue) in the healthy and WHS affected cerebellum, scale = 30 µm, inset = 15 µm. (F) Quantification of apoptotic astrocytes (GFAP and CL.C3 double positive) show a trend towards an increase in apoptosis in the WHS affected cerebellum, n = 3 hedgehogs/group, 2-tailed t-test, p = 0.05.

We also examined the astrocytes for cell death and observed a strong trend for increase in number of GFAP labelled cells co-expressing cleaved caspase 3 in the affected cerebellum (affected: 7.17 ± 2.6 X10^3^/mm^3^ vs healthy: 2.83 ± 0.74 X10^3^/mm^3^) (Fig. 5E), but it was barely significant (Fig. 5F). Overall, these results indicate that neuronal degeneration aided by astrocytic cell death is a major cause of the progressive paresis and paralysis in WHS.

## 4. Discussion

The presence of neuronal cell death along with extensive remyelination indicates that WHS is a neurodegenerative disease. WHS shares numerous characteristics with MS, a demyelinating neurodegenerative disease in humans. As in MS, WHS affected hedgehogs present with a variety of neurological signs depending on the location of CNS lesions (Diaz-Delgado et al., 2018; Graesser et al., 2006). Ataxia or incoordination are common presenting complaints, but other neurological signs like paraparesis, dysphagia, tremors, ascending paralysis, unilateral exophthalmos, and seizures have been reported (Graesser et al., 2006). The initial clinical signs have been described as relapsing and remitting, although this has not been thoroughly documented (Graesser et al., 2006). It is unknown at what point in the disease course demyelination and remyelination occur in hedgehogs affected with WHS. We have demonstrated that remyelination is sufficient to compensate for any oligodendroglial loss. Since WHS cannot be diagnosed antemortem, our current understanding of the disease is based on evaluation of the central nervous system in animals where neurological signs have already developed. Additionally, affected animals often display progressive neurological signs for weeks to months before euthanasia (Diaz-Delgado et al., 2018; Graesser et al., 2006). These limitations make it difficult to determine the existence of acute and chronic stages of WHS. In our study, all the affected hedgehogs presented with ataxia without any obvious relapsing and remitting disease course. Most WHS affected hedgehogs were tetraparetic but still ambulatory when euthanized for analysis at about 2 to 4 months after the initial onset of clinical signs.

A novel finding of this study is the loss of LFB staining with intact myelin proteins in the cerebellum of WHS affected hedgehogs (Fig. 1). LFB binds to phospholipids in the myelin sheath, which is composed of >70% lipids with the rest being myelin proteins (Pearse, 1955; Salthouse, 1962). The clinical signs and myelin changes in WHS closely resemble those in mice that cannot synthesize one of the major lipids, GalC and its sulfated derivative, sulfatide. These mice are able to form compact myelin with normal ultrastructural appearance. However, with age they develop progressive hindlimb paralysis and extensive vacuolation of the ventral region of the spinal cord with thinner myelin sheaths, but the levels of myelin proteins remain unchanged, similar to our findings in WHS (Coetzee et al., 1996). Reduction of myelin phospholipids without changes in the protein levels has also been observed in the diffusely abnormal white matter of MS brains, suggesting the possibility of a primary lipid abnormality in MS and WHS (Laule et al., 2013).

The ultrastructural findings in this study further extends our knowledge on the consequences of myelin vacuolation which leads to extensive demyelination and demonstrates the occurrence of remyelination and its extent, which was not reported in earlier studies. Axons of all diameters had thin myelin sheaths with high g-ratios, the hallmark of remyelination (Fig. 2, Fig. S3, Fig. S4) (Bunge et al., 1961; Duncan et al., 2017; Prineas and Connell, 1979). This extensive remyelination was likely due to regeneration of oligodendrocytes as evidenced by the brightly labelled CC1 expressing oligodendrocytes in the WHS affected cerebellum (Fig. 4). However, restoration of oligodendrocyte numbers could not compensate for the widespread neuronal cell death, one of the hallmarks of neurodegeneration that is also observed in primary progressive MS (Bramow et al., 2010; Lassmann et al., 2012; Manrique-Hoyos et al., 2012; Wilson et al., 2023). It is also possible that surviving adult oligodendrocytes participated in remyelination as shown in feline irradiated diet-induced demyelination (FIDID) (Duncan et al., 2009). Indeed, the myelin pathology in WHS strongly resembles that described in FIDID.

Unlike the relapsing remitting form of MS, there are very few therapeutic options for progressive forms of MS (Amezcua, 2022). Experimental autoimmune encephalitis is the most commonly used animal model for progressive MS, but it does not recapitulate every aspect of the human disease (Birmpili et al., 2022). Our results show that WHS closely resembles primary progressive MS with respect to the clinical course of the disease, evidence of extensive demyelination, remyelination, and neurodegeneration. However, the cause of WHS remains unknown and future studies directed towards understanding the etiology of this disease may help develop therapeutic strategies for progressive MS.

## Declaration of Competing Interest

JS is a co-inventor in the patent # US 9,248,128.

## Data availability

Data will be made available on request.

## Acknowledgements

Conceptualization (GAD, JS), experimental design (GAD, DZR, JS), data acquisition and regenerative analysis (DZR, JS, GAD, AK, MJR, AEP, BKA, IDD), writing (GAD, JS, IDD), funding (DZR, JS)

## Funding

This article was funded by the University of Wisconsin-Madison Stem Cell Medicine Center Postdoctoral Fellowship (DZR), NIH/NINDS grants R01NS119517 (JS) and R03NS126993 (JS), the Boespflug Myopathic Research Foundation (JS), and the University of Wisconsin – Madison Office of the Vice Chancellor for Research and Graduate Education with funding from the Wisconsin Alumni Research Foundation (JS).

## Appendix A. Supplementary data

**Table S1:** Primary antibodies used for immunofluorescence. List of primary antibodies along with the manufacturer information and dilutions used for immunofluorescence.

| **Name** | **Isotype** | **Catalog #** | **Manufacturer** | **Dilution** |
| --- | --- | --- | --- | --- |
| Olig2 | Ms<br>IgG2a | MABN50 | EMD Millipore | 1:200 |
| CC1 | Ms<br>IgG2b | OP80 | Calbiochem | 1:100 |
| MOG | Ms<br>IgG1 | Hybridoma | Gift from Alexander Gow | 1:100 |
| PLP1 | Rat<br>IgG | Hybridoma (clone AA3) | Gift from Alexander Gow | 1:50 |
| Beta III Tubulin | Ms<br>IgG2a | MAB1195 (Tuj1 clone) | Novus Biologicals | 1:100 |
| HuC/D | Rb IgG | Ab184267 | Abcam | 1:100 |
| Cleaved Caspase 3 | Rb IgG | 9661S | Cell Signaling | 1:200 |

**Figure S1:**
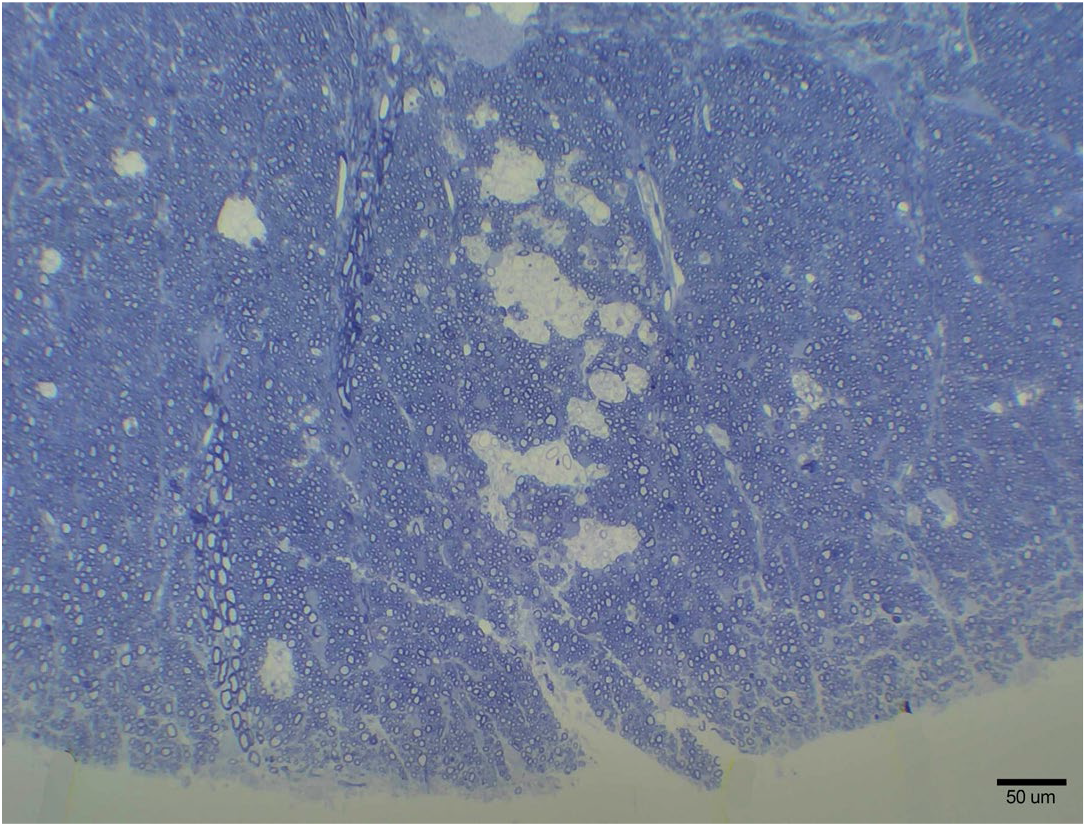
Scattered single vacuolated areas. In the youngest affected hedgehog, vacuolation was more limited than in the more severely affected animals but confirmed the diagnosis of WHS. Scale bar = 50 µm.

**Figure S2:**
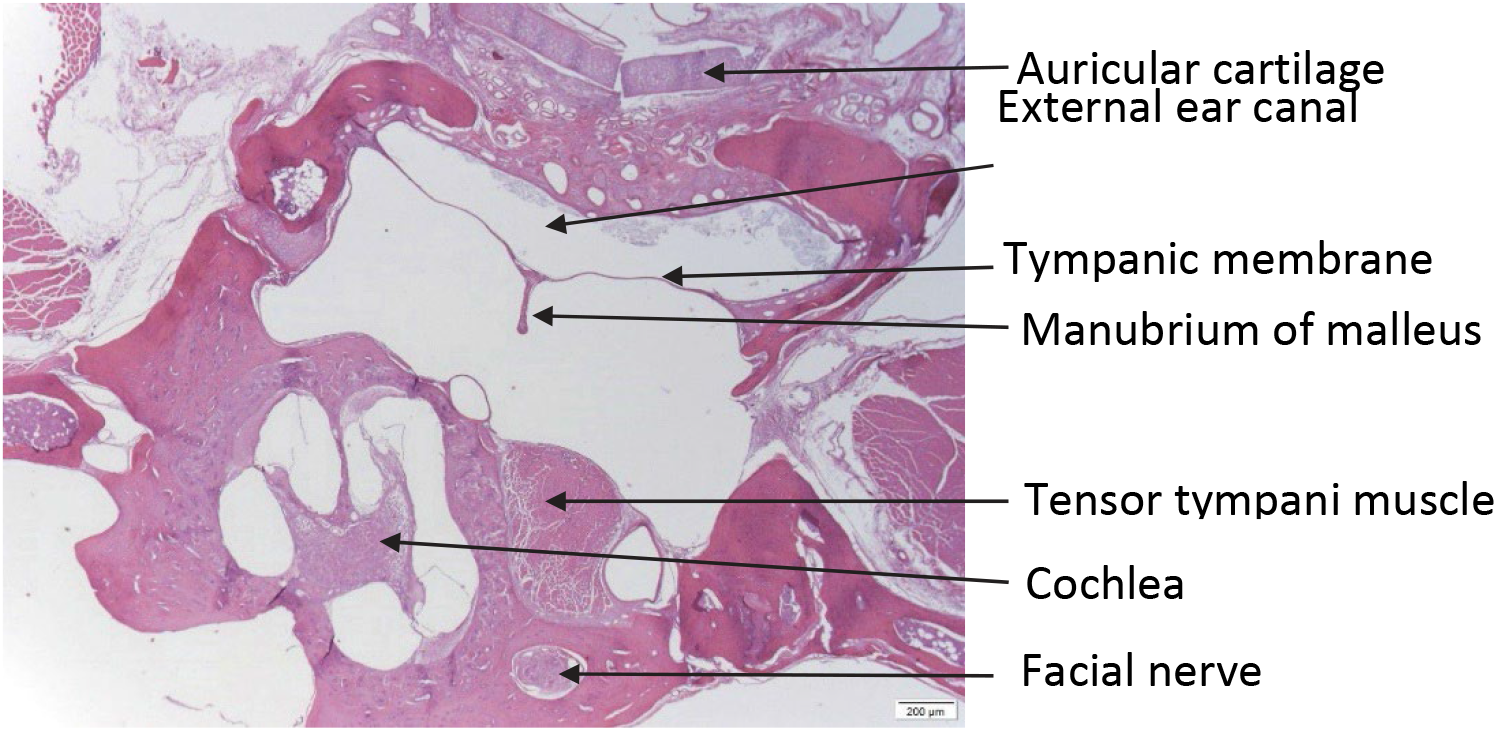
Normal bulla in WHS. Hematoxylin and eosin staining showed no abnormalities in the WHS affected bulla ruling out middle ear disease as a cause of ataxia.

**Figure S3:**
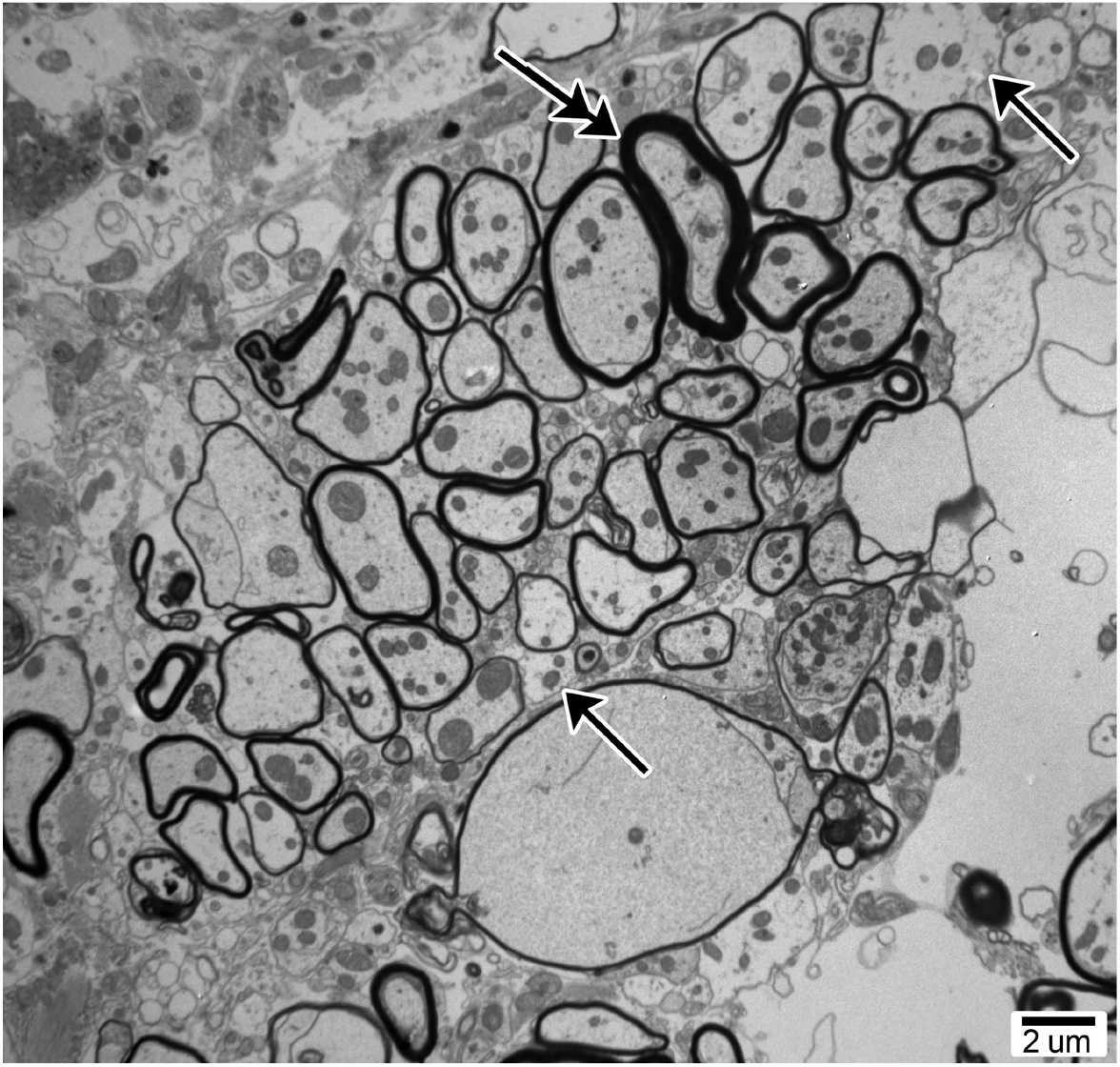
Contiguous areas of remyelination. While remyelinated axons more frequently scattered throughout the neuropil, occasional ‘clumps’ of remyelination were also seen, in addition to demyelinated axons (arrows) and normally myelinated axons (double arrow). Scale bar = 2 µm.

**Figure S4:**
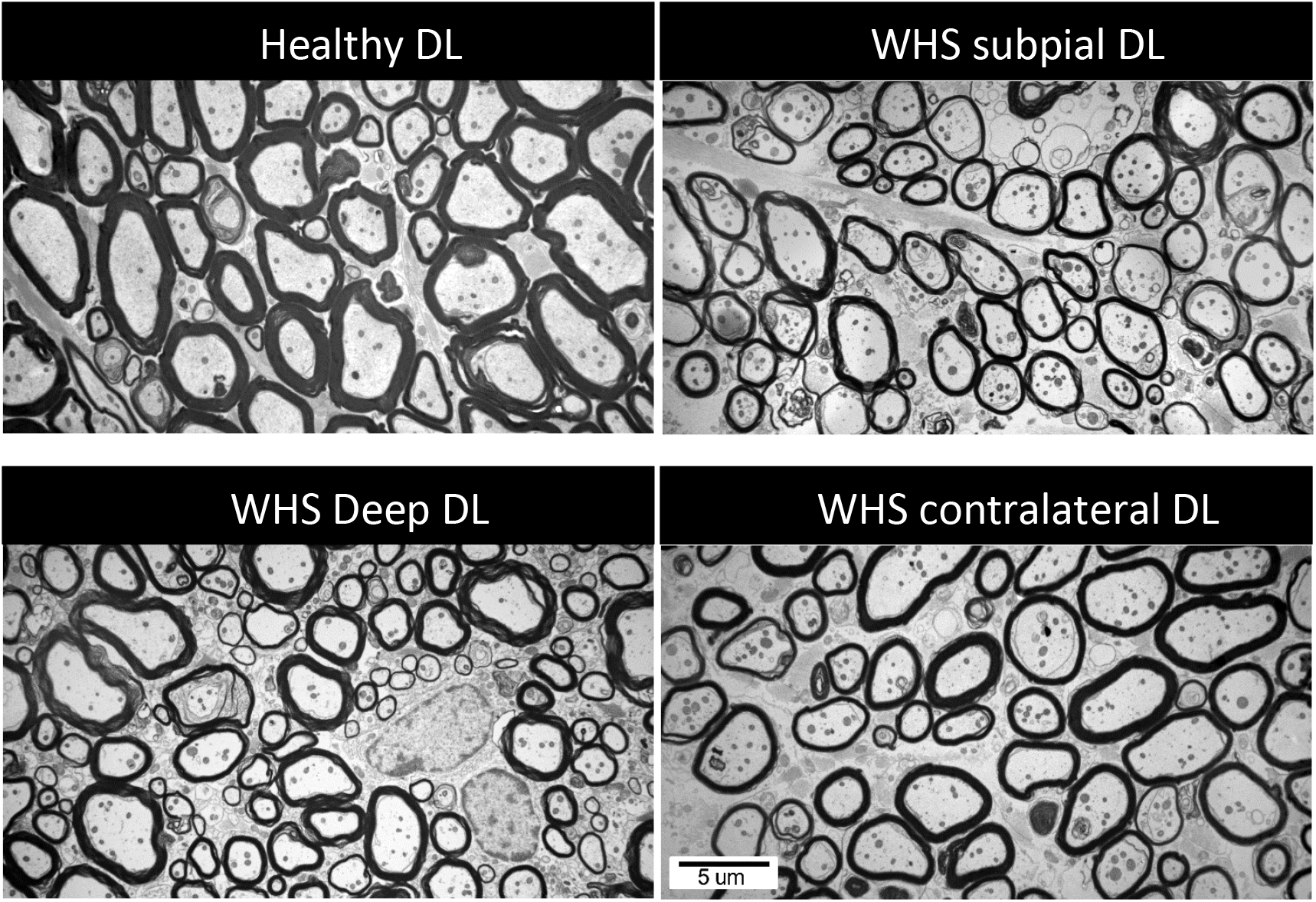
Ultrastructure of a young WHS affected cervical spinal cord. TEM images of the dorsolateral (DL) columns showed extensive remyelination in the right sub-pial area (upper right) but deep to this and close to the gray matter (lower left), axons of similar caliber have normal thickness myelin sheaths. On the opposite side (left), some axons have thin sheaths suggesting remyelination but not frequently as on the right side (lower right), Scale bar = 5 µm.

**Figure S5:**
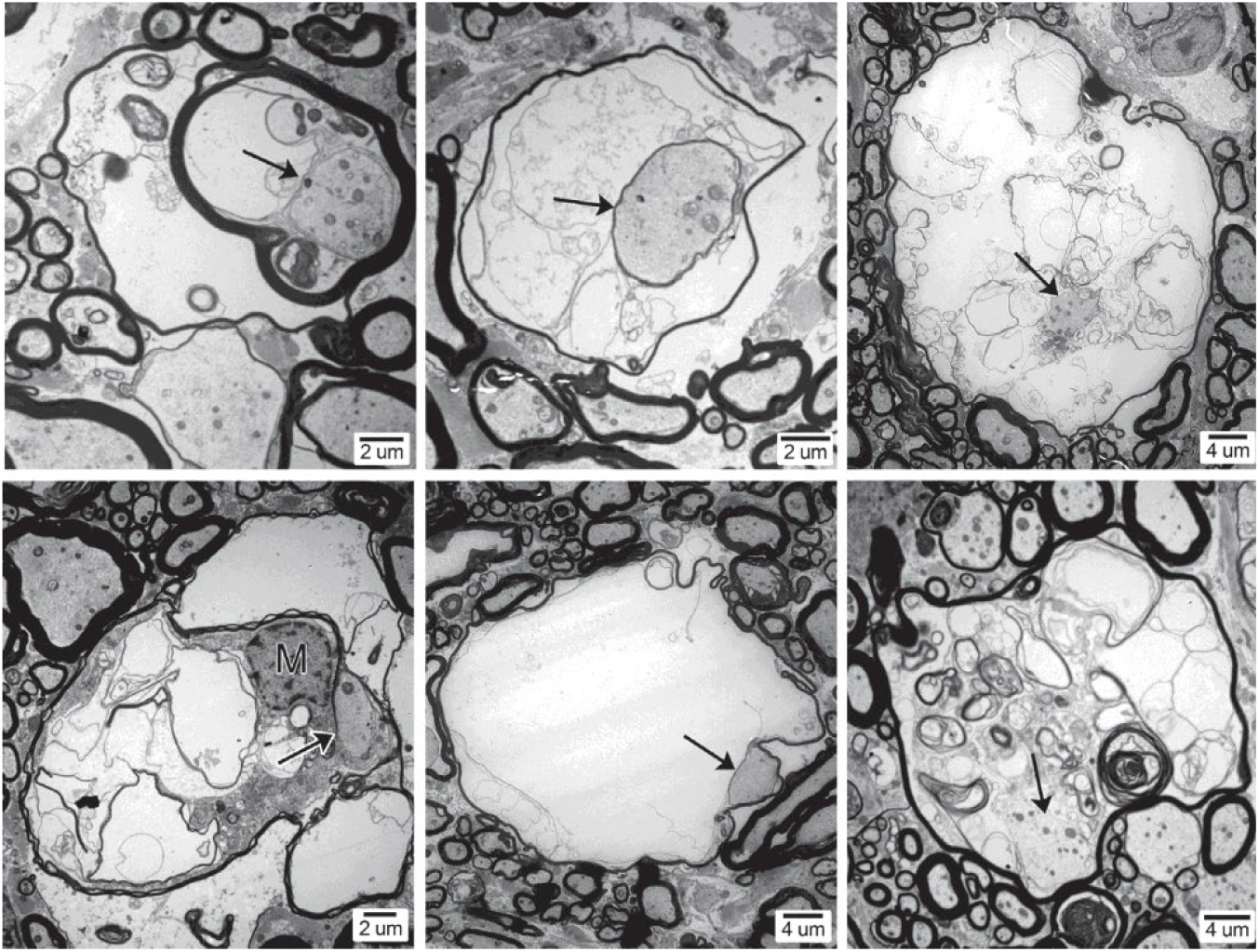
Ultrastructural evidence of degenerating myelin with intact axons. TEM images from ventral columns of WHS affected cervical spinal cords showed many vacuoles containing intact axons surrounded by degenerating myelin (arrows). In some cases, macrophages (M) were seen within the vacuole, ingesting the myelin debris. Scale bars = 2 and 4 µm.

**Figure S6:**
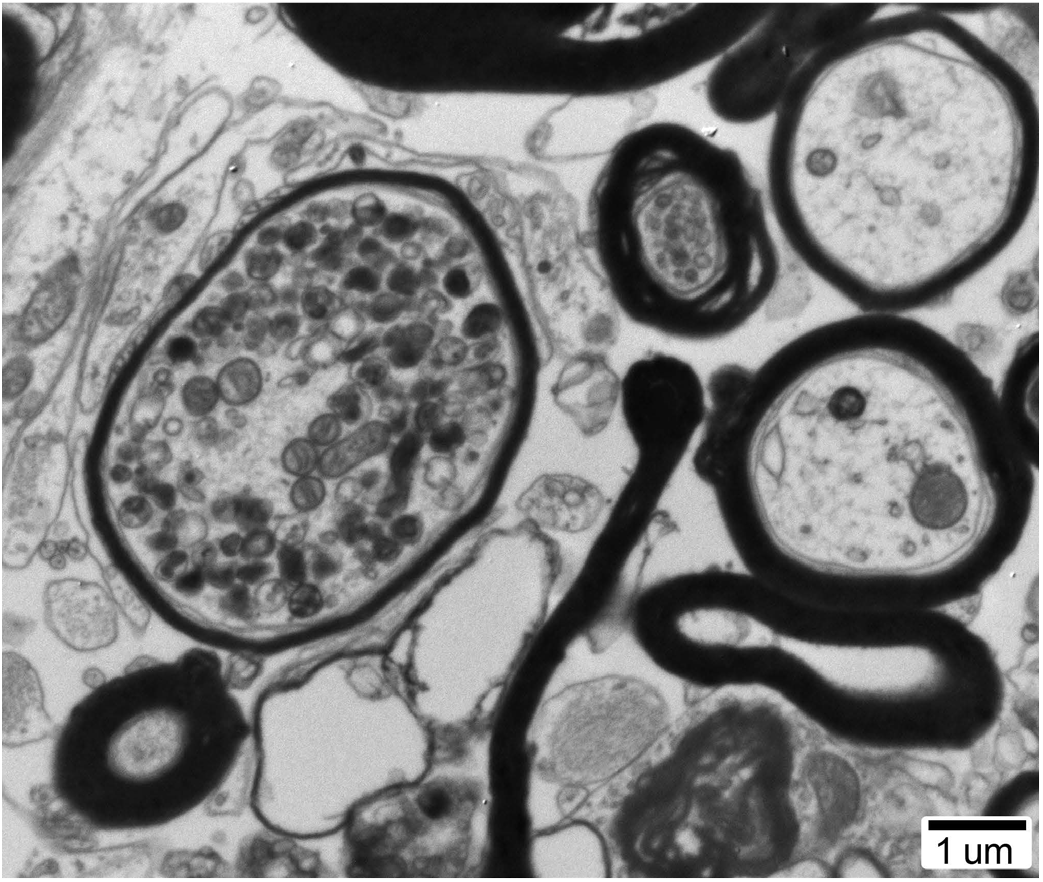
Axon degeneration. Rare axons with accumulated organelles, in this case, mitochondria, were noted. Scale bar = 1 µm.

**Figure S7:**
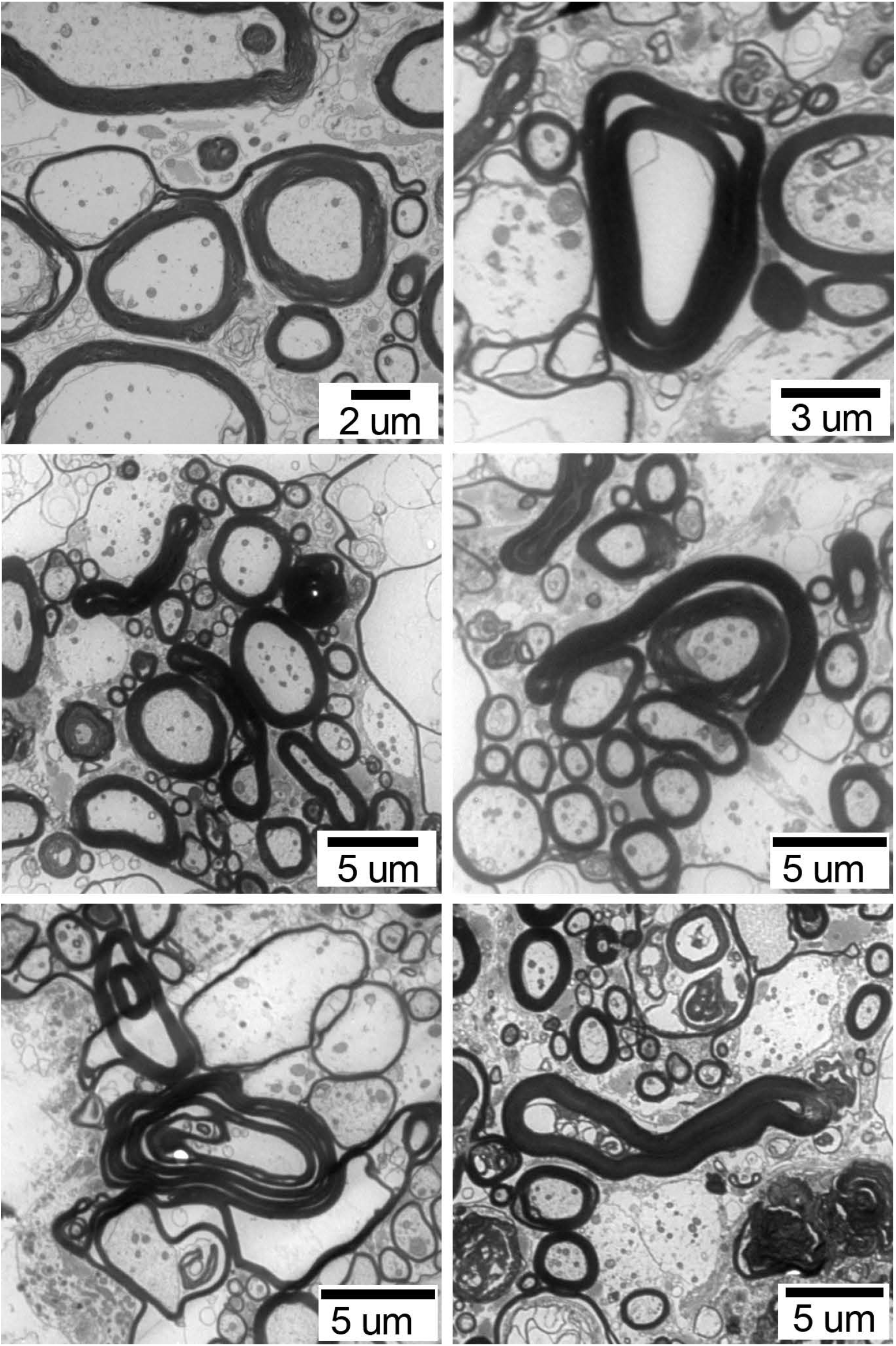
Redundant myelin. Many examples of redundant myelin sheaths, as seen in this figure, were found throughout the neuropil associated with myelin vacuoles and demyelinated and remyelinated axons. Scale bars = 2, 3 and 5 µm.

## Notes

### Competing Interest Statement

The authors have declared no competing interest.

## References

1. Amezcua, L. 2022. Progressive Multiple Sclerosis. Continuum (Minneap Minn). 28:1083–1103.

2. Bacmeister, C.M., R. Huang, L.A. Osso, M.A. Thornton, L. Conant, A.R. Chavez, A. Poleg-Polsky, and E.G. Hughes. 2022. Motor learning drives dynamic patterns of intermittent myelination on learning-activated axons. Nat Neurosci. 25:1300–1313.

3. Berg, C.C., G.A. Doss, and J. Guevar. 2021. Neurologic examination of healthy adult African pygmy hedgehogs (Atelerix albiventris). J Am Vet Med Assoc. 258:971–976.

4. Bin, J.M., S.N. Harris, and T.E. Kennedy. 2016. The oligodendrocyte-specific antibody ‘CC1’ binds Quaking 7. J Neurochem. 139:181–186.

5. Birmpili, D., I. Charmarke Askar, K. Bigaut, and D. Bagnard. 2022. The Translatability of Multiple Sclerosis Animal Models for Biomarkers Discovery and Their Clinical Use. Int J Mol Sci. 23.

6. Bramow, S., J.M. Frischer, H. Lassmann, N. Koch-Henriksen, C.F. Lucchinetti, P.S. Sorensen, and H. Laursen. 2010. Demyelination versus remyelination in progressive multiple sclerosis. Brain. 133:2983–2998.

7. Bunge, M.B., R.P. Bunge, and H. Ris. 1961. Ultrastructural study of remyelination in an experimental lesion in adult cat spinal cord. J Biophys Biochem Cytol. 10:67–94.

8. Carriel, V., A. Campos, M. Alaminos, S. Raimondo, and S. Geuna. 2017. Staining Methods for Normal and Regenerative Myelin in the Nervous System. Methods Mol Biol. 1560:207–218.

9. Chapman, T.W., G.E. Olveda, X. Bame, E. Pereira, and R.A. Hill. 2023. Oligodendrocyte death initiates synchronous remyelination to restore cortical myelin patterns in mice. Nat Neurosis. 26:555–569.

10. Clawson, E.D., D.Z. Radecki, and J. Samanta. 2023. Immunofluorescence assay for demyelination, remyelination, and proliferation in an acute cuprizone mouse model. STAR Protoc. 4:102072.

11. Coetzee, T., N. Fujita, J. Dupree, R. Shi, A. Blight, K. Suzuki, K. Suzuki, and B. Popko. 1996. Myelination in the absence of galactocerebroside and sulfatide: normal structure with abnormal function and regional instability. Cell. 86:209–219.

12. Diaz-Delgado, J., D.B. Whitley, R.W. Storts, J.J. Heatley, S. Hoppes, and B.F. Porter. 2018. The Pathology of Wobbly Hedgehog Syndrome. Vet Pathol. 55:711–718.

13. Djannatian, M., S. Radha, U. Weikert, S. Safaiyan, C. Wrede, C. Deichsel, G. Kislinger, A. Rhomberg, T. Ruhwedel, D.S. Campbell, T. van Ham, B. Schmid, J. Hegermann, W. Mobius, M. Schifferer, and M. Simons. 2023. Myelination generates aberrant ultrastructure that is resolved by microglia. J Cell Biol. 222.

14. Duncan, I.D., A. Brower, Y. Kondo, J.F. Curlee, Jr., and R.D. Schultz. 2009. Extensive remyelination of the CNS leads to functional recovery. Proc Natl Acad Sci U S A. 106:6832–6836.

15. Duncan, I.D., R.L. Marik, A.T. Broman, and M. Heidari. 2017. Thin myelin sheaths as the hallmark of remyelination persist over time and preserve axon function. Proc Natl Acad Sci U S A. 114:E9685–E9691.

16. Duncan, I.D., A.B. Radcliff, M. Heidari, G. Kidd, B.K. August, and L.A. Wierenga. 2018. The adult oligodendrocyte can participate in remyelination. Proc Natl Acad Sci U S A. 115:E11807–E11816.

17. Feldman, A.T., and D. Wolfe. 2014. Tissue processing and hematoxylin and eosin staining. Methods Mol Biol. 1180:31–43.

18. Franklin, R.J. 2002. Why does remyelination fail in multiple sclerosis? Nat Rev Neurosci. 3:705–714.

19. Franklin, R.J.M., and C. Ffrench-Constant. 2017. Regenerating CNS myelin – from mechanisms to experimental medicines. Nat Rev Neurosci. 18:753–769.

20. Franklin, R.J.M., J. Frisen, and D.A. Lyons. 2021. Revisiting remyelination: Towards a consensus on the regeneration of CNS myelin. Semin Cell Dev Biol. 116:3–9.

21. Frischer, J.M., S. Bramow, A. Dal-Bianco, C.F. Lucchinetti, H. Rauschka, M. Schmidbauer, H. Laursen, P.S. Sorensen, and H. Lassmann. 2009. The relation between inflammation and neurodegeneration in multiple sclerosis brains. Brain. 132:1175–1189.

22. Graesser, D., T.R. Spraker, P. Dressen, M.M. Garner, J.T. Raymond, G. Terwilliger, J. Kim, and J.A. Madri. 2006. Wobbly Hedgehog Syndrome in African Pygmy Hedgehogs (Atelerix spp.). Journal of Exotic Pet Medicine. 15:59–65.

23. Graves, J.S., K.M. Krysko, L.H. Hua, M. Absinta, R.J.M. Franklin, and B.M. Segal. 2022. Ageing and multiple sclerosis. The Lancet Neurology.

24. Grinspan, J., L. Wrabetz, and J. Kamholz. 1993. Oligodendrocyte maturation and myelin gene expression in PDGF-treated cultures from rat cerebral white matter. J Neurocytol. 22:322–333.

25. Kaiser, T., H.M. Allen, O. Kwon, B. Barak, J. Wang, Z. He, M. Jiang, and G. Feng. 2021. MyelTracer: A Semi-Automated Software for Myelin g-Ratio Quantification. eNeuro. 8.

26. Lassmann, H., J. van Horssen, and D. Mahad. 2012. Progressive multiple sclerosis: pathology and pathogenesis. Nat Rev Neurol. 8:647–656.

27. Laule, C., V. Pavlova, E. Leung, G. Zhao, A.L. MacKay, P. Kozlowski, A.L. Traboulsee, D.K. Li, and G.R. Moore. 2013. Diffusely abnormal white matter in multiple sclerosis: further histologic studies provide evidence for a primary lipid abnormality with neurodegeneration. J Neuropathol Exp Neurol. 72:42–52.

28. Madarame, H., K. Ogihara, M. Kimura, M. Nagai, T. Omatsu, H. Ochiai, and T. Mizutani. 2014. Detection of a pneumonia virus of mice (PVM) in an African hedgehog (Atelerix arbiventris) with suspected wobbly hedgehog syndrome (WHS). Vet Microbiol. 173:136–140.

29. Manrique-Hoyos, N., T. Jurgens, M. Gronborg, M. Kreutzfeldt, M. Schedensack, T. Kuhlmann, C. Schrick, W. Bruck, H. Urlaub, M. Simons, and D. Merkler. 2012. Late motor decline after accomplished remyelination: impact for progressive multiple sclerosis. Ann Neurol. 71:227–244.

30. Neely, S.A., J.M. Williamson, A. Klingseisen, L. Zoupi, J.J. Early, A. Williams, and D.A. Lyons. 2022. New oligodendrocytes exhibit more abundant and accurate myelin regeneration than those that survive demyelination. Nat Neurosci. 25:415–420.

31. Nikic, I., D. Merkler, C. Sorbara, M. Brinkoetter, M. Kreutzfeldt, F.M. Bareyre, W. Bruck, D. Bishop, T. Misgeld, and M. Kerschensteiner. 2011. A reversible form of axon damage in experimental autoimmune encephalomyelitis and multiple sclerosis. Nat Med. 17:495–499.

32. Palmer, A.C., W.F. Blakemore, R.J. Franklin, L.M. Frost, R.E. Gough, J.C. Lewis, D.F. Macdougall, M.T. O’Leary, and L.R. Stocker. 1998. Paralysis in hedgehogs (Erinaceus europaeus) associated with demyelination. Vet Rec. 143:550–552.

33. Pearse, A.G. 1955. Copper phthalocyanins as phospholipid stains. J Pathol Bacteriol. 70:554–557.

34. Peters, A. 2002. The effects of normal aging on myelin and nerve fibers: a review. J Neurocytol. 31:581–593.

35. Prineas, J.W., and F. Connell. 1979. Remyelination in multiple sclerosis. Ann Neurol. 5:22–31.

36. Revesz, T., D. Kidd, A.J. Thompson, R.O. Barnard, and W.I. McDonald. 1994. A comparison of the pathology of primary and secondary progressive multiple sclerosis. Brain. 117 (Pt 4):759–765.

37. Salthouse, T.N. 1962. A quantitative histochemical method for estimating phospholipids. Nature. 195:187–188.

38. Silva, G.F., A. Rema, S. Teixeira, M.D.A. Pires, M. Taulescu, and I. Amorim. 2022. Pathological Findings in African Pygmy Hedgehogs Admitted into a Portuguese Rehabilitation Center. Animals (Basel). 12.

39. Snaidero, N., M. Schifferer, A. Mezydlo, B. Zalc, M. Kerschensteiner, and T. Misgeld. 2020. Myelin replacement triggered by single-cell demyelination in mouse cortex. Nat Commun. 11:4901.

40. Solly, S.K., J.L. Thomas, M. Monge, C. Demerens, C. Lubetzki, M.V. Gardinier, J.M. Matthieu, and B. Zalc. 1996. Myelin/oligodendrocyte glycoprotein (MOG) expression is associated with myelin deposition. Glia. 18:39–48.

41. Trapp, B.D., and K.A. Nave. 2008. Multiple sclerosis: an immune or neurodegenerative disorder? Annu Rev Neurosci. 31:247–269.

42. Trapp, B.D., J. Peterson, R.M. Ransohoff, R. Rudick, S. Mork, and L. Bo. 1998. Axonal transection in the lesions of multiple sclerosis. N Engl J Med. 338:278–285.

43. Wilson, D.M., 3rd, M.R. Cookson, L. Van Den Bosch, H. Zetterberg, D.M. Holtzman, and I. Dewachter. 2023. Hallmarks of neurodegenerative diseases. Cell. 186:693–714.

44. Wu, H.Y., M.R. Dawson, R. Reynolds, and R.J. Hardy. 2001. Expression of QKI proteins and MAP1B identifies actively myelinating oligodendrocytes in adult rat brain. Mol Cell Neurosci. 17:292–302.

45. Yeung, M.S.Y., M. Djelloul, E. Steiner, S. Bernard, M. Salehpour, G. Possnert, L. Brundin, and J. Frisen. 2019. Dynamics of oligodendrocyte generation in multiple sclerosis. Nature. 566:538–542.

